# Graph neural representational learning of RNA secondary structures for predicting RNA-protein interactions

**DOI:** 10.1101/2020.02.11.931030

**Authors:** Zichao Yan, William L. Hamilton, Mathieu Blanchette

## Abstract

**Motivation:** RNA-protein interactions are key effectors of post-transcriptional regulation. Significant experimental and bioinformatics efforts have been expended on characterizing protein binding mechanisms on the molecular level, and on highlighting the sequence and structural traits of RNA that impact the binding specificity for different proteins. Yet our ability to predict these interactions *in silico* remains relatively poor.

**Results:** In this study, we introduce RPI-Net, a graph neural network approach for RNA-protein interaction prediction. RPI-Net learns and exploits a graph representation of RNA molecules, yielding significant performance gains over existing state-of-the-art approaches. We also introduce an approach to rectify particular type of sequence bias present in many CLIP-Seq data sets, and we show that correcting this bias is essential in order to learn meaningful predictors and properly evaluate their accuracy. Finally, we provide new approaches to interpret the trained models and extract simple, biologically-interpretable representations of the learned sequence and structural motifs.

**Availability:** Source code can be accessed at https://www.github.com/HarveyYan/RNAonGraph.

## 1 Introduction

RNA-protein interactions mediate all post-transcriptional regulatory events, including RNA splicing, capping, nuclear export, degradation, subcellular localization and translation (Stefl *et al.*, 2005). These regulatory processes are influenced by a diverse population of RNA binding proteins (RBPs), each having affinity for one or more specific RNA motifs. Defects in RBP-RNA interactions are implicated in a variety of neuromuscular disorders and possibly cancer (Lukong *et al.*, 2008; Xiong *et al.*, 2015). For this reason and many others, it is of crucial importance to adequately characterize the determinants of RBP binding specificity at the molecular level, including both sequence and structural motifs, as well as the broader impact of the sequence/structure context.

RBP binding has been shown to be determined by both the sequence and the structure of an RNA (Buckanovich and Darnell, 1997; Hackermuller *et al.*, 2005). For instance, Vts1p is an RBP that binds a certain sequence motif within a hairpin loop of RNA (Aviv *et al.*, 2006); therefore, prediction algorithms that do not consider secondary structures may fail to obtain optimal results. The difficulty of making use of RNA secondary structure in a machine learning context, however, is that it potentially requires modeling highly non-local interactions between pairs of nucleotides that could span hundreds or even thousands of bases in relatively long RNA transcripts.

Because of their nested structure, RNA secondary structures can be represented as simple strings (the so-called dot-bracket notation), and a number of recently proposed RBP binding prediction approaches combine these string-based structure representations with the nucleotide sequence information (Pan *et al.*, 2018; Kazan *et al.*, 2010; Cook *et al.*, 2017). However, the non-local interpretation of the dot-bracket notation makes it challenging for predominantly sequence-based models such as LSTMs to identify motifs that involve long-range interactions. A second fundamental challenge is to account for the stochasticity of RNA folding: instead of deterministically adopting their single most stable secondary structure, RNA molecules are constantly sampling from an ensemble of possible structures.

Therefore, in this study we introduce RPI-Net, a machine learning approach that aims at learning a more effective graph representation of the RNA secondary structures. Instead of manually extracting features with graph kernels (Maticzka *et al.*, 2014) or integrating other sources of hand-crafted features, RPI-Net is an end-to-end learning approach with graph neural networks (GNN) that learns directly at the molecular level from the sequence and structures of RNA. RPI-Net also captures the uncertainty involved in RNA folding by annotating edges with base-pairing probabilities.

RPI-Net is evaluated on a well-known RNA-protein interaction dataset from a collection of CLIP-Seq experiments (Hafner *et al.*, 2010; Licatalosi *et al.*, 2008; Konig *et al.*, 2010), originally assembled and curated by Maticzka *et al.* (2014). Compared to three state-of-the-art methods, RPI-Net improves prediction accuracy substantially on many RBPs. We also introduce new approaches based on Sundararajan *et al.* (2017) to interpret the trained models and reveal RBP sequence and structure binding preferences that often closely match experimentally derived motifs.

Finally, another contribution of our paper is to quantify the impact of and rectify a specific type of sequence bias present in certain CLIP-Seq datasets, originally identified by Ghanbari and Ohler (2019), which artificially inflated the reported prediction accuracy of several approaches published recently.

## 2 Related work

Our approach builds upon several computational biology frameworks that attempt to leverage RNA secondary structure information to predict RBP binding, and we extend this line of work by integrating recent advancements in graph neural networks (GNNs).

### Existing structure-aware bioinformatics tools

MEMERIS (Hiller *et al.*, 2006) is one of the earliest approaches to incorporating RNA secondary structure, though it has limited capacity and only models motifs in single-stranded (unpaired) areas of RNA. RNAcontext (Kazan *et al.*, 2010) later extends the structural context of RNA by annotating nucleotides with their structural state (stem, hairpin, bulge). This base annotation scheme is adopted in a series of more recent deep learning approaches (Zhang *et al.*, 2016; Pan *et al.*, 2018) that separately model the annotated secondary structures and primary sequence information in a multimodal fashion. Notably, Zhang *et al.* (2016) uses a multimodal deep belief network on the RNA tertiary structure as well as the sequence and secondary structure.

One key disadvantage of these multimodal approaches is that sequence and structure information are processed separately before being merged together. This can make it difficult to detect subtle relationships between sequential motifs and structural context. In addition, these approaches suffer from the limitation that they use a fixed sequential representation of the data, which makes higher-order interactions difficult to model and ignores the stochasticity of RNA folding.

There is also recent work, such as the Graphprot (Maticzka *et al.*, 2014) framework, that use a graph kernel to extract secondary structure features from an RNA graph, which simultaneously encode information about the nucleotide and its structural context. This approach is more effective at coupling sequence and secondary structure. However, the extremely high dimensionality of the extracted features can easily lead to overfitting.

### Graph neural networks

Recent advances of graph neural networks features two related genres. The first genre include a series of graph convolution-based models supported by spectral graph theory, which aim at learning approximate spectral filters defined on the graph Laplacian (Kipf and Welling, 2017). The other genre takes motivations from graphical models and graph isomorphism testing, and the operations are generalized as a form of differentiable message-passing between the nodes in a graph (Hamilton *et al.*, 2017). Methods of this type have been successfully applied to learn molecular fingerprints (Duvenaudt *et al.*, 2015) and to infer molecular properties (Gilmer *et al.*, 2017). In particular, adding recurrence to carry the state information when learning the node level embeddings has proved useful in the context of learning molecular graphs (Li *et al.*, 2016; Gilmer *et al.*, 2017), which motivates our choice of GNN architecture for RNA structures. Despite their prevalent usage in modeling molecular graph structure, applications of GNNs to RNA secondary structure have been scarce.

## 3 Methods

Figure 1(A) provides an overview of our RPI-Net approach. First, we fold RNA to obtain an ensemble of possible secondary structures and build a graph representation; then, we learn node level embeddings using a recurrent GNN and a bidirectional LSTM; finally, the node embeddings of an RNA are globally pooled along its spatial axis to make a final prediction of RNA-protein interaction for a given protein.

**Figure 1:**
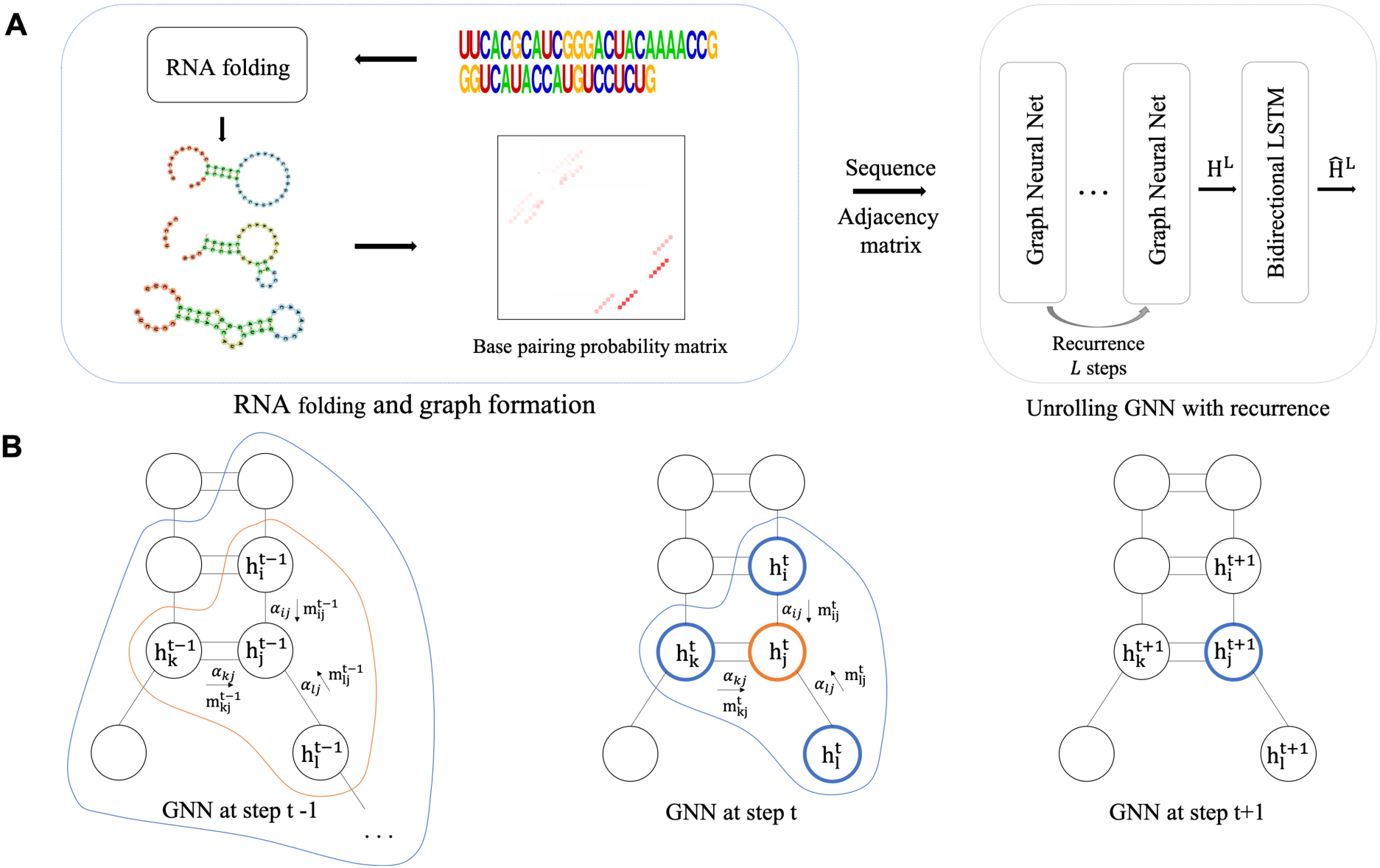
(a) Our graph neural network model takes in as input an adjacency matrix that specifies the base-pairing probabilities, as well as the one-hot encoded nucleotide sequence. The message passing layers are unrolled L steps using a long short-term memory model (LSTM) (Hochreiter and Schmidhuber, 1997), whose output is then fed to a bidirectional LSTM module to learn a more global nucleotide embedding. (b) Here we show an example of message passing to node j from its 2-hops neighbourhood. m_*ij*_ denotes the messages passed from node i to j, and to keep the notation uncluttered we have omitted its type and direction. *α*_*ij*_ is a scalar used for the base-pairing probability when m_*ij*_ is passed along a hydrogen bond, otherwise it is set to one. The node features associated to j at time step t is denoted as 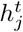, and it is updated with the information from its adjacent nodes according to Eq. 2 to Eq. 4.

### 3.1 RNA folding and sampling possible structures

To capture the dynamics and uncertainty of RNA secondary structure for a given sequence, RPI-Net uses an ensemble of possible structures sampled from the Boltzmann distribution, rather simply relying on the minimum free energy (MFE) structure. In this approach a secondary structure *s* associated with sequence *x* takes on probability 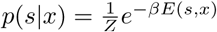, where *Z* is a normalizing constant (also known as the partition function), and *E*(*s, x*) is the free energy of *x* under structure *s*. The base-pairing probability for nucleotides at positions *i* and *j*, considering all possible secondary structures in a thermodynamic equilibrium, is then defined as:

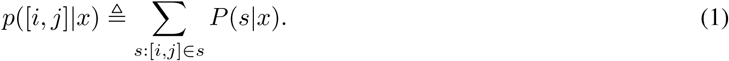

Even though the number of valid secondary structures increases exponentially as the sequence length increases, Eq. 1 can be calculated in time *O*(*n*^3^) using McCaskill’s algorithm (McCaskill, 1990), which is implemented in the RNAplfold package (Bernhart *et al.*, 2006). After running RNAplfold, we obtain a probabilistic adjacency matrix *A*_*n*×*n*_, where *A*_*i,j*_ = *p*([*i, j*]|*x*). These probabilistic annotations allow us to account for uncertainty in the RNA folding.

### 3.2 Learning nucleotide level embeddings

We start by giving a high-level overview of our approach, and provide full details later in the section. Our GNN is constructed based on the notion of neural message passing on an RNA graph formed by two types of bonds: covalent bonds linking consecutive nucleotides along the RNA backbone and base-pairing hydrogen bonds. Unlike many other applications of GNN, the probabilistically-weighted RNA graph is quite sparse, so in order to enable a more effective message passing, we choose to stack multiple GNN layers to obtain a sufficiently large receptive field. These layers share their weight parameters. This gives rise to a kind of recurrence resembling that of a Gated GNN model (Li *et al.*, 2016). In practice, we use a LSTM that treats the node embedding as a hidden state and the messages coming to each node as input. In addition, we compute the messages on the covalent bonds with a convolutional operation so that messages can be gathered from more distant nucleotides along the RNA backbone in a single GNN layer.

The core message passing function of RPI-Net takes three values as input: a probabilistic adjacency matrix *A*_*n*×*n*_; an embedding matrix 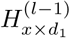, which contains a *d*_1_-dimensional embedding for each nucleotide and is updated by each message passing learning; and the LSTM cell memory *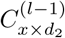*. The message propagation formula is then given by:

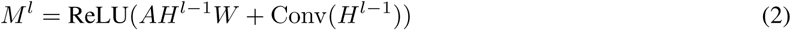

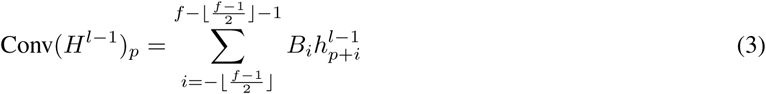

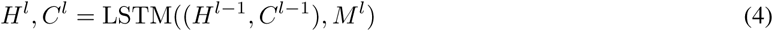

where ReLU is a leaky rectified linear unit (Maas *et al.*, 2013) and *W* and *B*_1_, …, *B*_*n*_ are trainable matrices. For the first layer of message passing (*l* = 1), we initialize *H*^0^ with an one-hot encoding of the nucleotide (i.e., {`A,U,C,G`}) and the cell memory *C*^0^ to zeros. The convolutional layer uses symmetric padding on both ends of the RNA sequence to keep its length constant. An example of message passing is given in Figure 1(B), where a nucleotide only receives covalent messages from its two immediately connected neighbors in addition to its base-pairs. We emphasize that—compared to a traditional GNN—we have generalized the message passing operation along the RNA backbone to a convolutional operation (Eq. 3). One can interpret Eq. 3 as a nucleotide positioned at index *p* receiving covalent messages from its neighborhood of size *f* (the receptive field of the convolution filter).

The total number of GNN layers is denoted by *L*, which is constrained to be a small number to avoid incurring a high computational overhead. To better account for long range dependencies between nucleotides, we feed *H*^*L*^ obtained at the final message passing step to a bidirectional LSTM module, which outputs a updated *Ĥ*^*L*^ as the nucleotide embeddings learned by our model.

### 3.3 Global RNA graph embedding

To obtain a global representation for the RNA as a whole, we finally use a Set2Set model (Vinyals *et al.*, 2016) to pool the individual nucleotide embeddings along the spatial axis of the RNA:

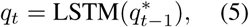

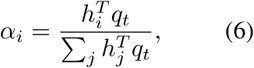

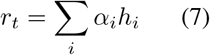

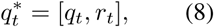

where each *h*_*i*_ is the row of *Ĥ*^*L*^ corresponding to nucleotide *i*. The Set2Set model is similar to the Seq2Seq (Sutskever *et al.*, 2014) model in that at each pooling step it uses attention to compare the decoder (a LSTM) hidden state to each nucleotide embedding (Eq. 6). The weighted sum of the nucleotide embeddings (Eq. 7) is concatenated with the hidden state (Eq. 8), which is then forwarded to the next pooling step as input to the decoder LSTM module (Eq. 5). The decoder is unrolled a fixed number of steps (*T*), and the concatenated vector 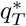 at the last step is used as an output from this module. The prediction is finally given by a softmax following a linear transformation on the output of the Set2Set model, which maps the dimensionality of 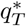 to two.

### 3.4 Hierarchical supervision

Our model is trained as follows. Each input sequence *s* in our training set is labeled with a “global” binary value *y*_*g*_(*s*) indicating whether or not it contains a binding site of the RBP of interest. We also associate to *s* a length *n* binary vector *y*_*l*_, where *y*_*l,i*_ indicates whether or not position *i* belongs to the viewpoint region. We train our model with a composite objective function combining a local and a global cross-entropy loss function. The global loss function is defined over an entire RNA sequence,

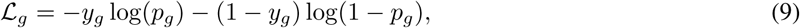

where *p*_rna_ is the predicted probability obtained as the output of the Set2Set module described in Section 3.3.In addition to this global loss, we can also define a local loss that operates directly on the individual nucleotide embeddings:

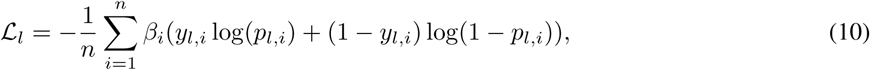

*β*_*i*_ is a positive scalar to weigh losses from different positions on a sequence.

Our complete minimization objective is given by the joint loss function

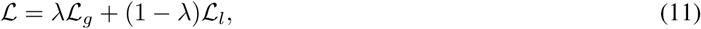

where λ is a mixing ratio controlling the importance of the two objectives.

Note that ℒ_*l*_ can provide a more fine-grained supervision signal than ℒ_*g*_, since we would like our model to discover the binding site primarily located within a relatively small viewpoint region, possibly aided by information gathered from the flanking regions.

However, ℒ_*l*_ is also arguably more challenging to optimize due to the highly disproportionate amount of nucleotides that are not within the marked viewpoint regions. To address this concern—which is similarly motivated in the field objection detection—we adopt a gradient harmonizing mechanism (GHM) (Li *et al.*, 2019) to mitigate the imbalance between positive and negative examples, or between easy and difficult examples, as well as to penalize outliers. GHM proceeds by estimating the gradient density GD(*g*_*i*_) for the gradient norm *g*_*i*_ on a per nucleotide basis. The gradient norm is taken from the loss with respect to the output of the model, indicating the difficulty of classifying that nucleotide. *β*_*i*_ is then simply given by 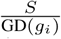. In practice, we also use a unit region approximation as well as the moving average statistics to compute GD(*g*_*i*_) more efficiently, as suggested by the authors of GHM.

### 3.5 Baseline predictors

To demonstrate the benefits of learning graph neural representations in a probabilistic ensemble setting, we also consider a baseline model that only looks at the RNA nucleotide sequence, by simply having a convolutional neural network (CNN) in place for GNN. Our baseline CNN model has two convolution layers with filters of length 10 and has the same number of hidden units as in GNN. Other components of the model are the same, such as the bidirectional LSTM layers and the Set2Set module. We apply the hierarchical objective function to both of our CNN and GNN models, which therefore are referred to as RPI-Net(CNN) and RPI-Net(GNN).

### 3.6 Sequence and secondary structure motif extraction

In biological applications, model interpretability (i.e. the ability to extract human-understandable, biologically-relevant information from a trained model) is nearly as important as prediction accuracy. In the context of studying RNA-protein interaction, this essentially means to uncover the sequence and secondary structural motifs that may describe an RBP’s binding affinity, as well as higher-level concepts such as cooperative binding.

#### Extracting sequence motifs from an RPI-Net(CNN) model

Integrated gradients (Sundararajan *et al.*, 2017) are an effective approach to assign an “attribution score” *Attr*(*i*) to each position *i* of a given input sequence *s*, measuring the extent to which the nucleotide at that position contributes to the entire sequence’s prediction score.

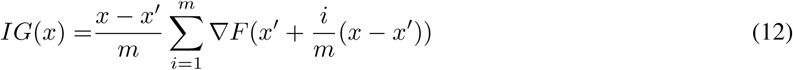

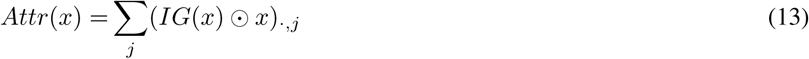

Since *F*, the logit from our neural network that makes the positive prediction for binding, is differentiable almost everywhere (Prop.1 in Sundararajan *et al.* (2017)), the following property holds: ∑_*i,j*_ *IG*(*x*)_*i,j*_ = *F*(*x*) − *F*(*x*′), where *x* is an one-hot encoded RNA sequence and *x*′ is a reference having the same length and feature dimension as *x. IG*(*x*) is a real-valued matrix where *i* indexes the position along the sequence and *j* indexes the encoded feature dimension at each position. Therefore, each entry in *IG*(*x*) records the contribution of that position in the sequence to *F*(*x*), and for the purpose of revealing attributions of the positive predictions of binding sites, *x* is selected from the positive RNA examples and *x*′ is simply fixed to zeros.

Since each position in *x* is encoded with only one nucleotide, we use *Attr*(*x*) from Eq. 13 as the final result of the integrated gradients, which is a vector that only retains the contribution that exists in the original sequence. Note that we use ⊙ to denote element-wise multiplication.

To identify portions of sequence *x* of highest relevance to the prediction, and considering that RBPs tend to recognize short 5-10 nt motifs, we assign to each position *i* the sum of the attribution scores of the *K* = 10 positions centered at *i*.

We identified the highest-scoring *K*-mer from each of the 2000 positive sequences with the highest prediction scores, and then produced a multiple alignments of those *K*-mers using *clustalw2* (Larkin *et al.*, 2007), to eventually derive a position weight matrices (PWM) profile, which can be visualized using *WebLogo* (Crooks *et al.*, 2004).

#### Extracting sequence and structural motifs from RPI-Net(GNN) models

The input to a graph neural network, unlike the pure CNN-based approach, is a sequence-structure pair. During training, the structure is an adjacency matrix filled with equilibrium base-pairing probabilities. At the stage of secondary structural motif extraction, whose purpose is to characterize the structural preference of the sequence binding motif, we only study one structure at a time, sampled from RNAsubopt (Lorenz *et al.*, 2011).

We note that we do not directly compute integrated gradients of the adjacency matrix. Once a secondary structure is sampled, we consider its adjacency matrix as fixed weight parameters of the network, and compute integrated gradients to the nucleotide sequence as usual. The secondary structure is annotated with *forgi* (Kerpedjiev *et al.*, 2015) which associates each position in the sequence with a structural element such as dangling start (F), dangling end (T), hairpin loop (H), internal loop (I), multi-loop (M) and stem (S). Therefore, for each sequence-structure pair, we extract a sequence *k*-mer using the aforementioned procedure, along with its structural annotation which is also a *k*-mer. The same multiple sequence alignment step is applied to the sequence *k*-mer, with gaps inserted in the structural *k*-mer at the same position as its corresponding sequence *k*-mer.

Note that given an identical sequence paired with different secondary structures, *Attr*(*x*) should be different for these two structures and a higher score would suggest the sequence *k*-mer is better recognized by the model due to its structural context. Therefore, to adequately characterize the structural motif, we only consider the sequence and structural *k*-mers with high attribution scores, otherwise the structural motif may become less meaningful because of the sheer quantities of random structural *k*-mers. For this purpose, we sample 2000 structures from each sequence (including the flanking regions to the viewpoint), and rank all the *k*-mers from each sequence-structure pair based on the attribution scores. The top ranked *k*-mers are then selected to build the sequence and structural motifs.

Due to the consideration of running time, we have only used the top 100 positive sequences based on the prediction scores, which in the end gives us 200,000 sequence 10-mers and their corresponding structural 10-mers. At the multiple sequence alignment step, out of these 200,000 10-mers, we only align the top 6000 sequence 10-mers with the highest attribution scores. Then gaps are inserted to the structural 10-mers at the same position as in their corresponding sequence 10-mers.

### 3.7 Debiasing CLIP-Seq data

Finally, before turning to our empirical results, we discuss a important data bias present in many CLIP-seq datasets and our methodology to rectify this bias. Certain variants of the CLIP-Seq protocol, such as PAR-CLIP (Hafner *et al.*, 2010) and HITS-CLIP (Licatalosi *et al.*, 2008), use particular RNA cleavage enzymes such as RNase T1 to separate a portion of RNA that contains the binding site from the original transcript, resulting in a snippet that can be later sequenced and mapped to the original genome. The issues related to these enzymes are that they favour a certain pattern of cleavage, such as cleaving after Guanine. Therefore, in many positive examples the RNA-seq reads that contain RBP binding sites would be preceded by Guanine and end as well with Guanine.

We adopt the viewpoint terminology originally introduced in Maticzka *et al.* (2014), referring to an approximate region centered in an RNA sequence, relative to the flanking regions extended in both upward (5’UTR) and downward (3’UTR) directions. For a positive RNA sequence, the viewpoint is an actual RNA-seq read that contains a binding site. In negative examples, viewpoints would be randomly shuffled unbound sites. Therefore, in dataset obtained with particular CLIP-Seq protocols such as PAR-CLIP and HITS-CLIP, viewpoints in the positive examples usually start with Guanine and end with Guanine. Such a pattern, however, does not appear in the negative examples, as shown in Figure 2. Notably, iCLIP (Konig *et al.*, 2010) as another variant of CLIP-Seq does not seem to exhibit this type of bias.

**Figure 2:**
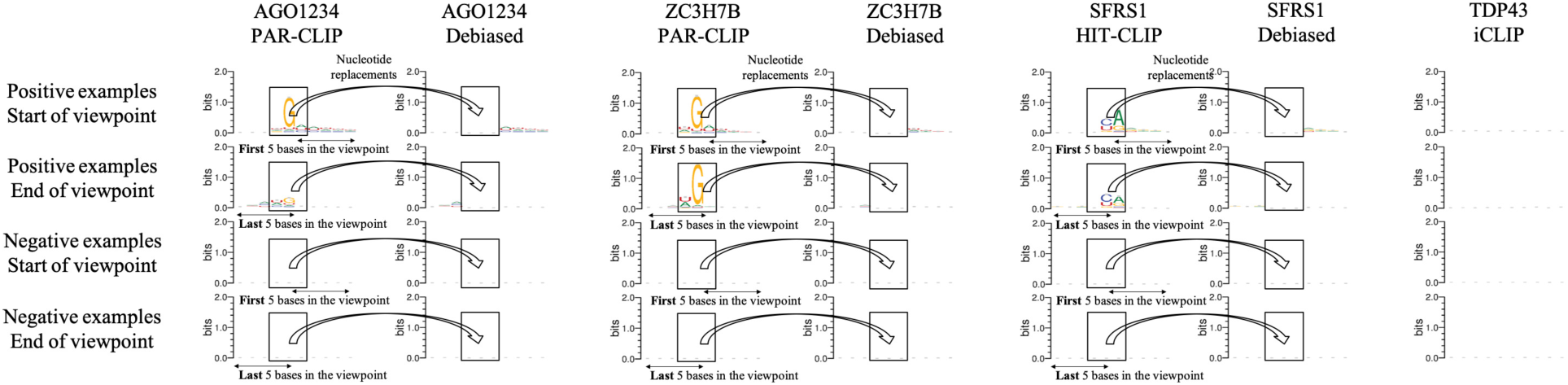
Positive examples from the PAR-CLIP dataset can be biased at the beginning and at the end of the viewpoint regions, emitting an unusually high frequency of Guanine (Ghanbari and Ohler, 2019) and some correlated residuals, which can be revealed by aligning the viewpoint borders of all positive examples. Such pattern, however, does not seem to exist in the negative examples, e.g. in the two RBP binding dataset for AGO1-4 and ZC3H7B. It can be further qualitatively verified that all PAR-CLIP data have suffered from this type of bias, and the complete set of evidence can be found in the Figure S1. HITS-CLIP is another protocol that can also be affected, with analogous but arguably milder border artefacts. SFRS1 is provided as an example for this family of protocols. Other dataset obtained with iCLIP, for example TDP43, does not appear to have this type of bias at all.

This cleavage bias is critical, since machine learning algorithms that base their decisions on features extracted in relatively close vicinity to the viewpoint borders will very likely end up learning border motifs that strongly favour Guanine, rather than actual RBP binding motifs. To address the above concern, we propose a simple yet effective approach to rectify the bias by replacing the nucleotides at positions that emit this border pattern with uniformly drawn random nucleotides. We replace the nucleotides inside a window of size 3 centered at the Guanine preceding the viewpoint, as well as those within another window of size 3 centered at the end of the viewpoint, as shown in Figure 2. The same replacement strategy is used for all affected RBP datasets. We employ this strategy for both positive examples and negative examples so that a machine learning predictor will not merely learn to distinguish altered (or in other words, unnatural) nucleotide sequences from the original ones.

After this nucleotide replacement procedure, we create a debiased version of the dataset where the viewpoint border artefacts have been largely removed, although it can be observed that for some RBP datasets such a border bias may extend more than 3 bases in both directions, e.g. EWSR1, ELAVL1 (C) and QKI in Figure S1. We select windows of size 3 to guarantee that Guanine and its two nearby nucleotides are always replaced. Larger window size would unavoidably degrade the quality of RNAs by changing their underlying semantics.

With this debiased dataset, we can move on to evaluate a machine learning algorithm more fairly since it would be pushed to learn meaningful RBP binding motifs instead of simply exploiting the border statistics.

## 4 Results

### 4.1 Datasets

We evaluate our approach on a well-known benchmarking RBP binding dataset originally curated by Maticzka *et al.* (2014), which features 24 human RBP binding experiments derived from two earlier studies (Anders *et al.*, 2012; Xue *et al.*, 2009) based on different CLIP-Seq protocols (PAR-CLIP, iCLIP and HITS-CLIP). Each positive sequence consists of a “viewpoint” region of 12-75 nt, which is the region identified by the experiment, flanked on each side with 150 nt (when possible), for a total length ranging from 38 to 375. The flanking regions provide the necessary context to study the region’s secondary structure and potentially locate binding sites for cooperative binding. Negative examples are obtained by choosing at random viewpoint-sized portions of human transcripts, ensuring they do not overlap positive examples for that RBP, and then extending them by 150 nt. Most data sets include 8,000-50,000 positive examples, and roughly equally many negative examples (Table S2).

Out of the 24 RBP datasets, 16 are obtained with PAR-CLIP and are thus subject to the bias described in Section 3.7. Another 4 are obtained with HITS-CLIP that are also similarly biased, except for the PTB dataset. The four Iclip datasets do not exhibit any visible sequence bias. Our de-biasing procedure (see Methods) was applied to all PAR-CLIP and HITS-CLIP (except PTB) data sets (both positive and negative examples), which effectively eliminated this unwanted signal (Figure S1). The same debiased datasets were used to train and evaluate all models.

### 4.2 Performance Comparisons

#### PAR-CLIP bias impacts previous machine learning methods

In Table S3 we present evidence that two representative algorithms, iDeepE (Pan and Shen, 2018) and GraphProt (Maticzka *et al.*, 2014), are indeed affected by the PAR-CLIP sequence bias.

iDeepE is an ensemble model combining a global and a local convolution neural network to predict RBP binding from RNA sequence only. Here we demonstrate that when an iDeepE model fitted with the original CLIP-Seq data is used to predict on the debiased data, its accuracy declines substantially, especially for RBPs such as CAPRIN1, AGO1-4, IGF2BP123, MOV10 and ZC3H7B, which from Figure S1 are also those where the PAR-CLIP sequence bias is strongest. This suggests that iDeepE’s previously reported accuracy may be inflated by this experimental bias. Indeed, iDeepE’s local convolution neural network can be particularly susceptible to this type of bias. Nonetheless, iDeepE remains a strong predictor on debiased data for many RBPs.

GraphProt, which bases its predictions on features mainly extracted from the viewpoint and surrounding nucleotides, is also affected by this bias, but to lesser extent. In particular, its performance on AGO1-4 and ZC3H7B is strongly impacted. This can be corroborated with evidence from GraphProt’s original paper, which presents the learnt motifs for these two RBPs (Maticzka *et al.* (2014); Figure 7), showing a close resemblance between GraphProt’s learned motifs and the border motifs we show in Figure 2. Still, based on results presented in Table S3, GraphProt is more resilient to the PAR-CLIP artefact than its deep learning competitor in almost every RBP dataset. This suggests that the high expressivity of deep neural networks could be a double-edged sword, should the training data contain any bias that can potentially be exploited by the model. Therefore, it is imperative that a model should only see debiased CLIP-Seq data during training, otherwise it can be affected by the bias, unless special training strategy is used to shield the model from exposure to the viewpoint border statistics, which we leave for future works.

#### State-of-the-art performance is achieved by RPI-Net

Here, we compare our approach to three popular RBP-binding predictors: iDeepE, GraphProt and mDBN.

mDBN is another deep learning approach that encodes RNA sequence and secondary structures using replicated softmax topic model, complemented by a tertiary structural profile. The training of mDBN first undergoes a series of pre-training iterations as a deep belief net, and is then trained by stochastic gradient descent as a feed-forward neural net. We found ourselves unable to re-evaluate mDBN using the debiased data, because the tool is no longer maintained and lacks documentation; hence the performance reported here is based on the author’s own evaluation on the un-debiased data set.

Table 1 and Figure 3 report 10-fold cross-validation results on debiased data for iDeepE, GraphProt, and our approach. Detailed hyperparameters used to evaluate our models can be found in Table S1. Both our CNN or GNN approaches consistently outperform the three existing predictors in terms of AUROC values, with average AUROC gains of 2.89%, 7.29%, and 3.92% compared to mDBN+ (non-debiased data), GraphProt, and iDeepE respectively (p-value < 0.001 for all comparisons, calculated for Wilcoxon signed-rank test (Wilcoxon, 1946) with alternative hypothesis that AUROC values of RPI-Net(GNN) are greater). In all 24 datasets, the AUROC values obtained by our two predictors exceed that of GraphProt and iDeepE on the same debiased data, while mDBN+ (evaluated on non-debiased data) comes out as a winner for 4 RBPs.

**Table 1:** ROC AUC scores from 24 RBP binding experiments, based on 10-fold cross-validation. Underlined RBP indicates that the original dataset is biased and a debiased version is used to train the models. AUC scores that are within 1% of the best score obtained for a given RBP are shown in bold.

| RBP | mDBN+<br>(original) | GraphProt<br>(debiased) | iDeepE<br>(debiased) | RPI-Net<br>(CNN)<br>(debiased) | RPI-Net<br>(GNN)<br>(debiased) |
| --- | --- | --- | --- | --- | --- |
| <u>ALKBH5</u> | 0.686 | 0.671 | 0.632 | 0.714 | **0.724** |
| <u>C17ORF85</u> | 0.817 | 0.774 | 0.770 | **0.848** | **0.844** |
| <u>C22ORF28</u> | 0.792 | 0.729 | 0.786 | **0.840** | **0.849** |
| <u>CAPRIN1</u> | 0.834 | 0.790 | 0.786 | 0.843 | **0.869** |
| Ago2 | 0.809 | 0.755 | 0.851 | **0.888** | 0.877 |
| <u>ELAVL1(H)</u> | **0.966** | 0.953 | **0.970** | **0.973** | **0.971** |
| <u>SFRS1</u> | **0.931** | 0.894 | 0.909 | **0.936** | **0.941** |
| HNRNPC | 0.962 | 0.952 | **0.978** | **0.985** | **0.986** |
| TDP43 | 0.876 | 0.874 | 0.941 | **0.961** | **0.959** |
| TIA1 | 0.891 | 0.861 | 0.941 | **0.964** | **0.963** |
| TIAL1 | 0.870 | 0.833 | 0.933 | **0.960** | **0.959** |
| Ago1-4 | 0.881 | 0.819 | 0.868 | **0.920** | **0.927** |
| <u>ELAVL1(B)</u> | **0.961** | 0.930 | 0.944 | **0.963** | **0.964** |
| <u>ELAVL1(A)</u> | **0.966** | 0.954 | **0.969** | **0.972** | **0.968** |
| <u>EWSR1</u> | **0.966** | 0.926 | 0.946 | **0.964** | **0.967** |
| <u>FUS</u> | **0.980** | 0.945 | 0.968 | **0.978** | **0.980** |
| <u>ELAVL1(C)</u> | **0.994** | 0.983 | **0.990** | **0.995** | **0.995** |
| <u>IGF2BP1-3</u> | 0.879 | 0.854 | 0.843 | **0.904** | **0.912** |
| <u>MOV10</u> | 0.854 | 0.757 | 0.818 | 0.846 | **0.875** |
| <u>PUM2</u> | **0.971** | 0.951 | 0.940 | **0.968** | **0.972** |
| <u>QKI</u> | **0.983** | 0.952 | 0.958 | **0.976** | **0.977** |
| <u>TAF15</u> | **0.983** | 0.966 | 0.967 | **0.982** | **0.981** |
| PTB | **0.983** | 0.937 | 0.944 | 0.958 | 0.958 |
| <u>ZC3H7B</u> | 0.796 | 0.675 | 0.764 | 0.793 | **0.838** |

**Figure 3:**
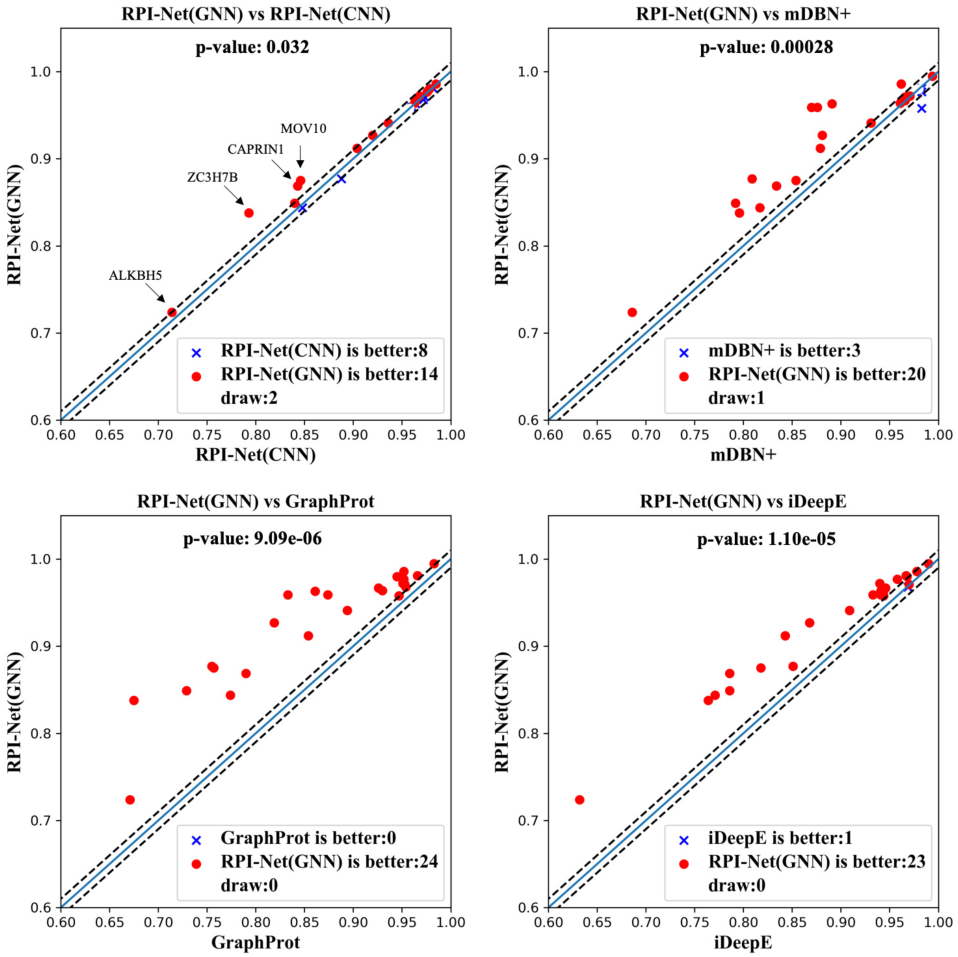
Comparison of RPI-Net(GNN) ROC AUC values to previous models and RPI-Net(CNN). The dotted lines correspond to a 1% difference. The p-value is calculated for the Wilcoxon signed-rank test with the alternative hypothesis that AUC scores of RPI-Net(GNN) are greater.

#### Incorporating secondary structure improves performance

The RPI-Net(CNN) and RPI-NET(GNN) approaches perform comparably on many datasets, but the latter achieves superior results for 14 RBPs (against only 8 for which the CNN is better). This notably includes four RBPs (ALKBH5, CAPRIN1, MOV10 and ZC3H7B) for which the AUROC gain exceeds 1% using the GNN, highlighting the utility of incorporating secondary structures in the learning process.

### 4.3 Motif visualizations

#### Sequence motifs from RPI-Net(CNN) model

To interpret the trained models, we developed a summarization approach based on integrated gradients and sequence motif analysis (Section 3.6). The integrated gradient approach attributes a score to each position in a sequence, measuring the extent to which the model’s prediction depends (positively or negatively) on the nucleotide at that position. Figure 4(A) shows the integrated gradients values for four positive examples for different RBPs, based on models trained on the non-debiased (top) and debiased (bottom) data. The former models assigns very high attribution scores to the “G” nucleotide near the viewpoint’s boundaries, and comparatively lower scores elsewhere. However, with models trained on debiased data, this border artefacts disappears and meaningful protein binding motifs emerge within the viewpoints. Notably, regions with high attribution scores tend to resemble known motifs for those RBPs (panel B, left column).

**Figure 4:**
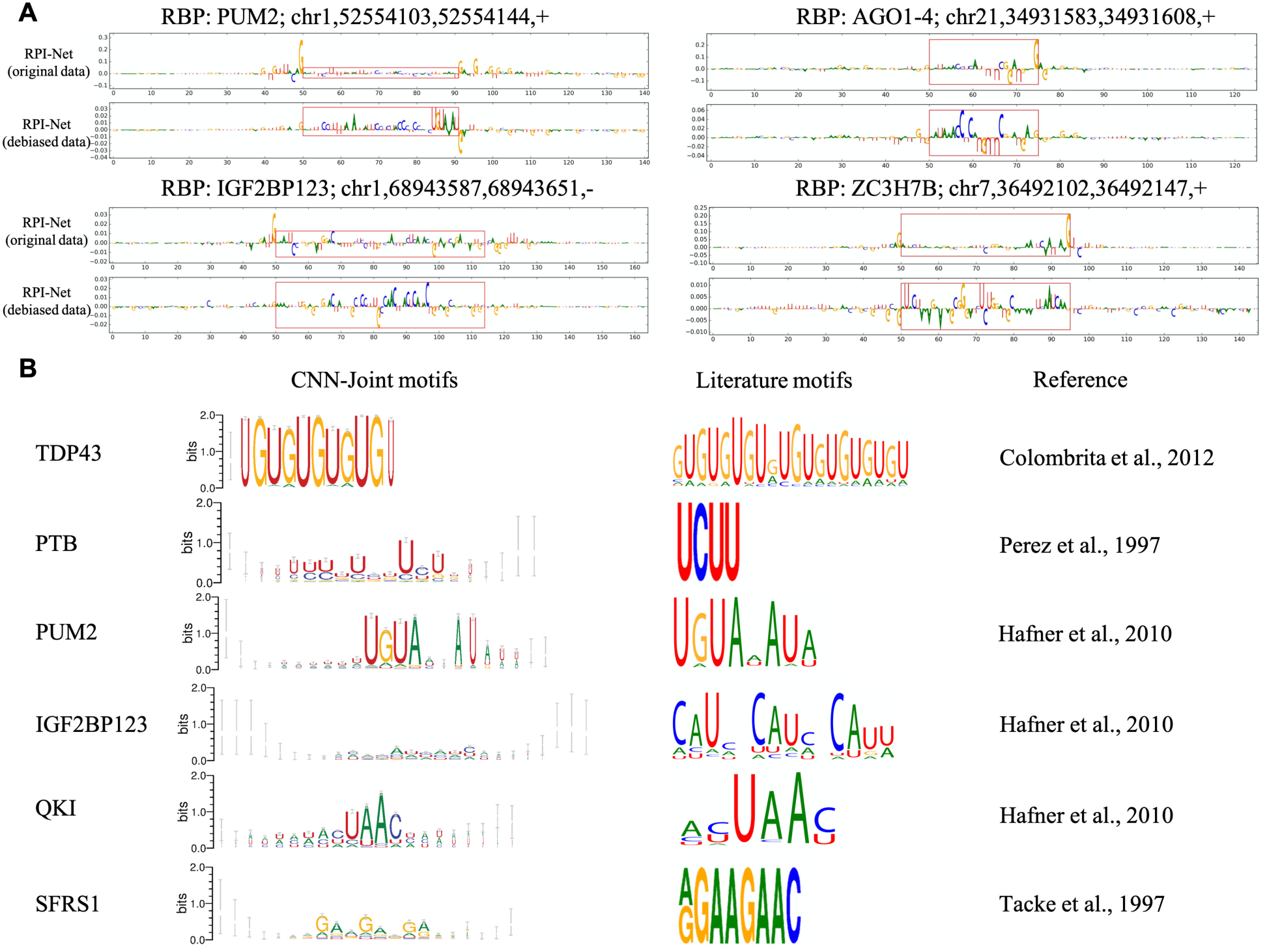
Integrated gradient maps and sequence motifs obtained using RPI-Net(CNN). (A) Integrated gradient values for four positive examples, revealing that debiasing the CLIP-Seq data is crucial for RPI-Net(CNN) to learn meaningful protein binding hypothesis. When RPI-Net is trained on the original (biased) datasets, the border artefacts will receive the highest attribution of the model’s prediction. Trained on debiased datasets, the border artefacts is eliminated and meaningful protein binding pattern are revealed inside the viewpoint region (enclosed red rectangles). (B) shows six sequence logos extracted from RPI-Net(CNN) models, compared to the known motifs for those RBPs. The widths of characters become smaller as more gaps are inserted to that position. Our sequence motifs have shown a strong agreement to the literature motifs that are experimentally verified. Image credits: The literature motif images are taken from Maticzka *et al.* (2014), Figure 5.

While attribution scores can highlight relevant portions of individual input sequences, they need to be combined to obtain a more global representation of the sequence motifs sought by our models. We developed an approach based on alignment and summarization of substrings with high attribution scores (see Methods), which results in a more classical sequence logo for each RBP. Figure 4 (B) shows the motifs obtained for six RBPs for the RPI-Net(CNN) model; the complete list of motifs for all RBP is available in Figure S2.

Many of motifs found have shown a strong agreement with experimentally-determined motifs such as TBP43, PTB, PUM2, QKI and SFRS1 (Hafner *et al.*, 2010; Tacke *et al.*, 1997; Colombrita *et al.*, 2012; Perez *et al.*, 1997). In general, the majority of our sequence motifs shown in Figure S2 conform well to the prior biological observations, including A-U rich patterns for EWSR1, TAF15 and FUS, and U-track for HNRNPC and TIA1. There are also less successful cases such as IGF2BP1-3 where the motif pattern is less conspicuous, potentially due to the uncertainty of the model about the binding mechanism of that RBP. This also tends to be the cases where the RPI-Net(CNN) model makes less accurate predictions (e.g. C22ORF28, AGO1-4, MOV10, C17ORF85, ZC3H7B and CAPRIN1). This suggests that the quality of motifs extracted from the model is dictated by the model’s capability of making accurate predictions for an RBP.

#### Identification of secondary sequence motifs

In addition to the sequence motifs obtained by multiple sequence alignment, we investigated an alternate approach using motif discovery program MEME (Bailey *et al.*, 2015) to identify enriched motifs within the 10-mers identified using our integrated gradients approach. A potential advantage of tools like MEME is that they can discover multiple motifs, some of which may be mediating cooperative binding with other RBPs. Results are shown in Figure S3. For several RBPs (e.g. TIAL1, SFRS1 and AGO1-4), we can observe more than one motif with very low e-values that may be attributed to different RBPs. Notably, most of the motifs found here were not detected by directly running MEME on the entire viewpoint regions, showing that our approach was instrumental in highlighting key portions of those sequences.

#### Sequence and structural motifs from RPI-Net(GNN) model

Extracting motifs from the GNN model is more challenging because they involve a combination of sequence and secondary structure. Applying the approach described in Methods, we obtained paired sequence-structure logos for each RBP. Figure 5 shows the motifs obtained for the RBPs for which the GNN approach yielded significantly improved accuracy compared to the CNN (see Figure S4 for the full results).

**Figure 5:**
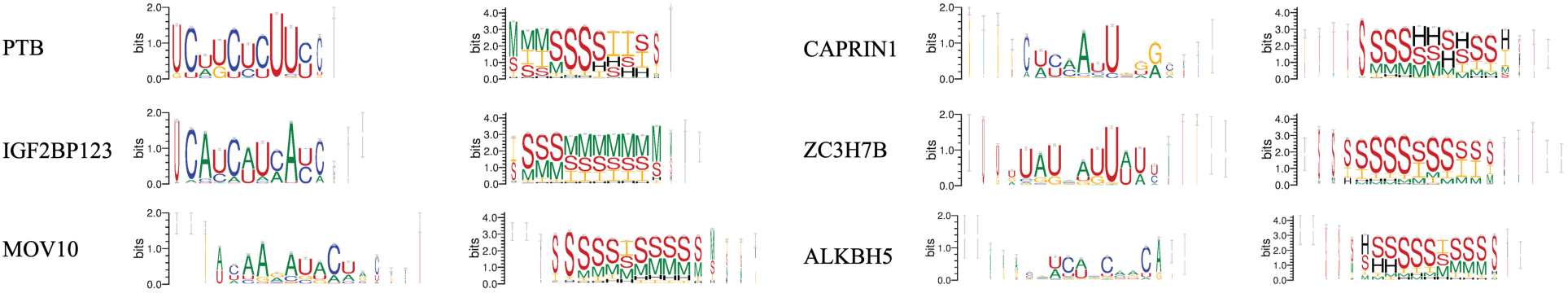
Six representative sequence and secondary structure motifs extracted from RPI-Net(GNN), including the four RBPs (MOV10, CAPRIN1, ZC3H7B and ALKBH5) where RPI-Net(GNN) outperformed the RPI-Net(CNN). Note how the motifs identified for PTB and IGF2BP123 resemble the literature motifs in Figure 4. The structural motifs are composed with a letter set of F (dangling start), T (dangling end), H (hairpin loop), I (internal loop), M (multi-loop) and S (stem).

Despite having a limited quantity of RNA sequences, we can still reveal some remarkable binding hypothesis learnt by our GNN model. The PTB binding motif shows a clear “UCUU” repeat that is almost absent in its CNN counterpart. The structural motif of PTB also highlights segments of unpaired regions separated by a double-stranded region, which suggests a stem-loop structural component to the PTB binding motif. The IGF2BP1-3 motif also demonstrates a clear “CAU” repeat that strongly agrees with the literature, which also tends to adopt a structural context of stem-loops. The structural motifs for MOV10, CAPRIN1, ZC3H7B and ALKBH5, suggest that these sequence motifs are mostly located in double-stranded areas, except for CAPRIN1, which tends to have a small segment of a hairpin loop.

## 5 Discussion and conclusion

In this study we introduced a novel approach to model RNA secondary structures with graph neural networks (GNNs), which shows important performance gains over the previous methods and improves on our own powerful CNN baseline. We can also interpret the meaningful protein binding hypothesis learnt by our models by extracting sequence and structural motifs. We also propose a strategy to eliminate CLIP-Seq border artefacts, which enable our approach to learn meaningful protein binding hypotheses.

Despite the significant performance gains and meaningful biological interpretations provided by our GNN as well as CNN models, our approach can be still improved in many aspects to better integrate RNA secondary structure information. For one, our approach is not taking advantage of the nested nature of RNA secondary structure, which could be naturally represented by a junction tree or hypergraph (Jin *et al.*, 2018). For instance, one could consider aggregating structurally adjacent nucleotides within the same double-stranded stem or unpaired loop region into a single node in a hypergraph. This hierarchical pooling strategy could enable our model to learn more meaningful graph level embeddings and to make faster inference, since the pooling operations interleaving the GNN layers would reduce the dimensionality of the graph while also refining the higher level node embeddings with more hierarchical information.

Our work also demonstrates that the removal of border artefacts is crucial for an end-to-end learning system to learn non-trivial protein binding hypothesis. An interesting alternative to our de-biasing technique is the one introduced by Ghanbari and Ohler (2019), who formulate the RBP binding prediction problem as a multi-class classification, aiming to simultaneously predict the binding of all RBPs in a collection of CLIP-seq data sets, which combines multiple biased RBP dataset into one. If all datasets are equally affected by sequence biases introduced by the experimental protocol, then this bias is uninformative for the prediction task and should not significantly affect the training. However, our experience is that different PAR-CLIP datasets exhibit biases of different strengths, which would be problematic for this approach; the criterion of grouping different protein dataset should be based on the similarity of their border emission patterns. A proper quantitative guidance for merging different dataset has yet to be defined.

One other aspect that calls for additional improvement is related to the secondary structural motifs extracted from our GNN models, which are limited in the sense that they are merely presented in a one dimensional format, although the inputs to the GNN is represented as a full two dimensional base-pairing probability matrix. To obtain secondary structural motifs in the two dimensional space, a subgraph alignment technique may be developed to obtain a probabilistic profile over the structural preference around the binding sites.

Finally, a transcript’s secondary structure may allow binding sites for cooperating RBPs to come in close physical proximity (despite being distant along the linear sequence) as suggested in Figure S5, and hence allow stabilizing protein-protein interactions. A potential benefit of the GNN approach that remains to be explored is its ability to amalgamate sequence signal information across the entire structure in a manner that may be more biologically relevant than what a CNN can achieve.

## Acknowledgements

We thank Eric Lécuyer, Jérôme Waldispühl, Yanlin Zhang and Faizy Ahsan for helpful discussions. We also thank Yue Li and Carlos G. Oliver for reviewing our manuscript. We are also thankful for the GPU allocations on Compute Canada.

