## Supplementary file for "Graph neural representational learning of RNA secondary structures for predicting RNA-protein interactions"

### A Hyperparameters

Table S1: Common hyperparameters for all the experiments.

| | |
|---|---|
| window size/span (RNAplfold) | 150 |
| probability cutoff (RNAplfold) | 1e-4 |
| no lonely basepair (RNAplfold) | True |
| hidden units (in the graph message passing layer and the Set2Set module) | 32 |
| convolutional filter length | 10 |
| $L$ (LSTM unroll steps) | 10 |
| $T$ (Set2Set pooling steps) | 10 |
| dropout ratio | 0.2 |
| learning rate | 2e-4 |
| batch size | 128 |
| $\lambda$ (hierarchical loss mixing ratio) | 0.95 |
| optimizer | AMSGrad |
| validation ratio (to select the best model) | 0.1 |
| epochs | 400 |

### B Dataset information

Table S2: Dataset size and average sequence length (nt).

| RBP | Dataset size | Positive examples | Negative examples | Length |
|---|---|---|---|---|
| C22ORF28 | 18,505 | 9,369 | 9,136 | 330±20 |
| CAPRIN1 | 16,041 | 8,140 | 7,901 | 334±23 |
| Ago2 | 92,346 | 48,095 | 44,251 | 335±23 |
| ELAVL1 (H) | 17,031 | 8,595 | 8,436 | 337±20 |
| SFRS1 | 36,633 | 19,438 | 17,195 | 333±32 |
| HNRNPC | 41,266 | 21,472 | 19,794 | 338±13 |
| TDP43 | 167,110 | 92,031 | 75,079 | 347±15 |
| TIA1 | 34,184 | 18,049 | 16,135 | 340±28 |
| TIAL1 | 78,984 | 42,332 | 36,652 | 344±24 |
| Ago1–4 | 68,212 | 36,902 | 31,310 | 327±27 |
| ELAVL1 (B) | 51,249 | 27,275 | 23,974 | 332±21 |
| ELAVL1 (A) | 18,747 | 9,464 | 9,283 | 337±19 |
| EWSR1 | 31,012 | 16,292 | 14,72 | 326±17 |
| FUS | 66,061 | 34,581 | 31,480 | 324±13 |
| ELAVL1 (C)/HUR | 238,888 | 125,202 | 113,686 | 323±14 |
| IGF2BP1-3 | 15,377 | 8,539 | 6,838 | 353±25 |
| MOV10 | 26,780 | 13,793 | 12,987 | 330±24 |
| PUM2 | 17,343 | 9,116 | 8,227 | 323±36 |
| QKI | 19418 | 10,276 | 9,142 | 327±27 |
| TAF15 | 13,904 | 7,298 | 6,606 | 322±19 |
| PTB | 88,274 | 44,574 | 43,700 | 325±14 |
| ZC3H7B | 40,890 | 20,962 | 20,018 | 341±19 |

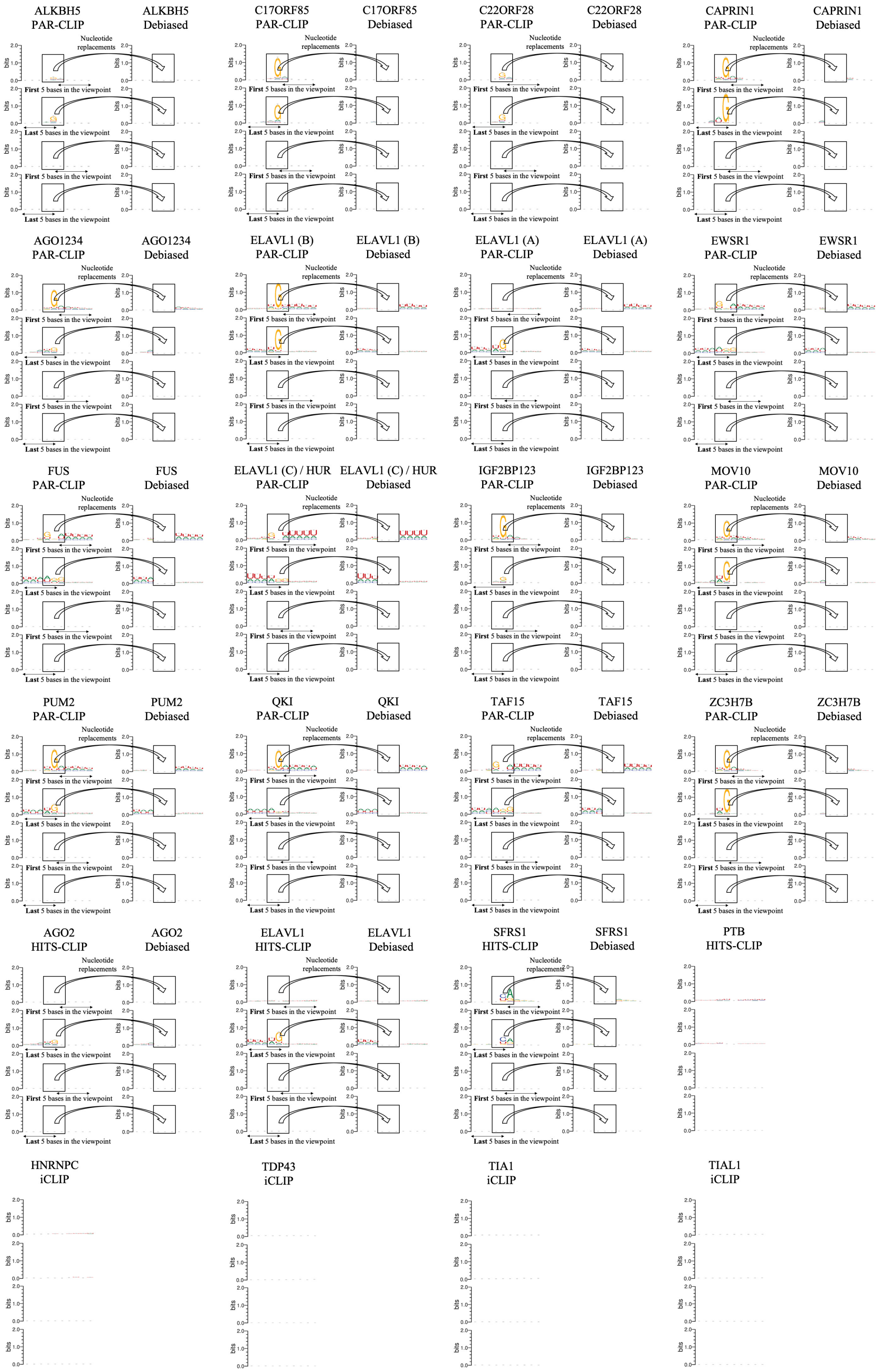


Figure S1: The debiasing scheme for all dataset except for PTB, HNRNPC, TDP43, TIA1 and TIAL1 which are not biased. For each RBP dataset, the first and second rows denote the viewpoint border nucleotides replacement scheme for the positive examples. The same procedures are applied to the negative examples, as shown in the third and fourth rows.

#### C PAR-CLIP bias impacts previous machine learning methods

Table S3: This table quantifies the impact of the PAR-CLIP data bias on iDeepE and GraphProt. Models are trained with the original CLIP-Seq data, and then evaluated on either the same data or the debiased data (in all cases, using 10-fold cross-validation). The difference between the model's performance on the two types of test sets measures the sensitivity of the models to the PAR-CLIP artefact.

| | iDeepE | | | GraphProt | | |
|---|---|---|---|---|---|---|
| RBP | original | debiased | diff | original | debiased | diff |
| ALKBH5 | 0.665 | 0.621 | -6.62% | 0.681 | 0.664 | -2.50% |
| C17ORF85 | 0.815 | 0.742 | -8.96% | 0.781 | 0.762 | -2.43% |
| C22ORF28 | 0.809 | 0.765 | -5.44% | 0.742 | 0.725 | -2.29% |
| CAPRIN1 | 0.896 | 0.687 | -23.33% | 0.842 | 0.787 | -6.53% |
| Ago2 | 0.881 | 0.827 | -6.13% | 0.758 | 0.762 | +0.53% |
| ELAVL1(H) | 0.975 | 0.968 | -0.72% | 0.954 | 0.960 | +0.63% |
| SFRS1 | 0.935 | 0.892 | -4.60% | 0.899 | 0.898 | -0.11% |
| HNRNPC | 0.978 | - | - | 0.952 | - | - |
| TDP43 | 0.941 | - | - | 0.874 | - | - |
| TIA1 | 0.941 | - | - | 0.861 | - | - |
| TIAL1 | 0.933 | - | - | 0.833 | - | - |
| Ago1–4 | 0.927 | 0.759 | -18.12% | 0.861 | 0.790 | -8.25% |
| ELAVL1(B) | 0.975 | 0.896 | -8.10% | 0.935 | 0.944 | +0.95% |
| ELAVL1(A) | 0.975 | 0.967 | -0.82% | 0.955 | 0.961 | +0.63% |
| EWSR1 | 0.968 | 0.927 | -4.24% | 0.934 | 0.935 | +0.11% |
| FUS | 0.981 | 0.968 | -1.33% | 0.952 | 0.954 | +0.21% |
| ELAVL1(C) | 0.993 | 0.985 | -0.81% | 0.989 | 0.977 | -1.21% |
| IGF2BP1-3 | 0.922 | 0.751 | -18.55% | 0.882 | 0.857 | -2.83% |
| MOV10 | 0.906 | 0.742 | -18.10% | 0.766 | 0.767 | +0.13% |
| PUM2 | 0.959 | 0.917 | -4.38% | 0.958 | 0.953 | -0.52% |
| QKI | 0.967 | 0.958 | -0.93% | 0.955 | 0.955 | 0.00% |
| TAF15 | 0.976 | 0.967 | -0.92% | 0.969 | 0.968 | -0.10% |
| PTB | 0.944 | - | - | 0.937 | - | - |
| ZC3H7B | 0.902 | 0.645 | -28.49% | 0.740 | 0.653 | -11.76% |

#### D All RPI-Net(CNN) motifs

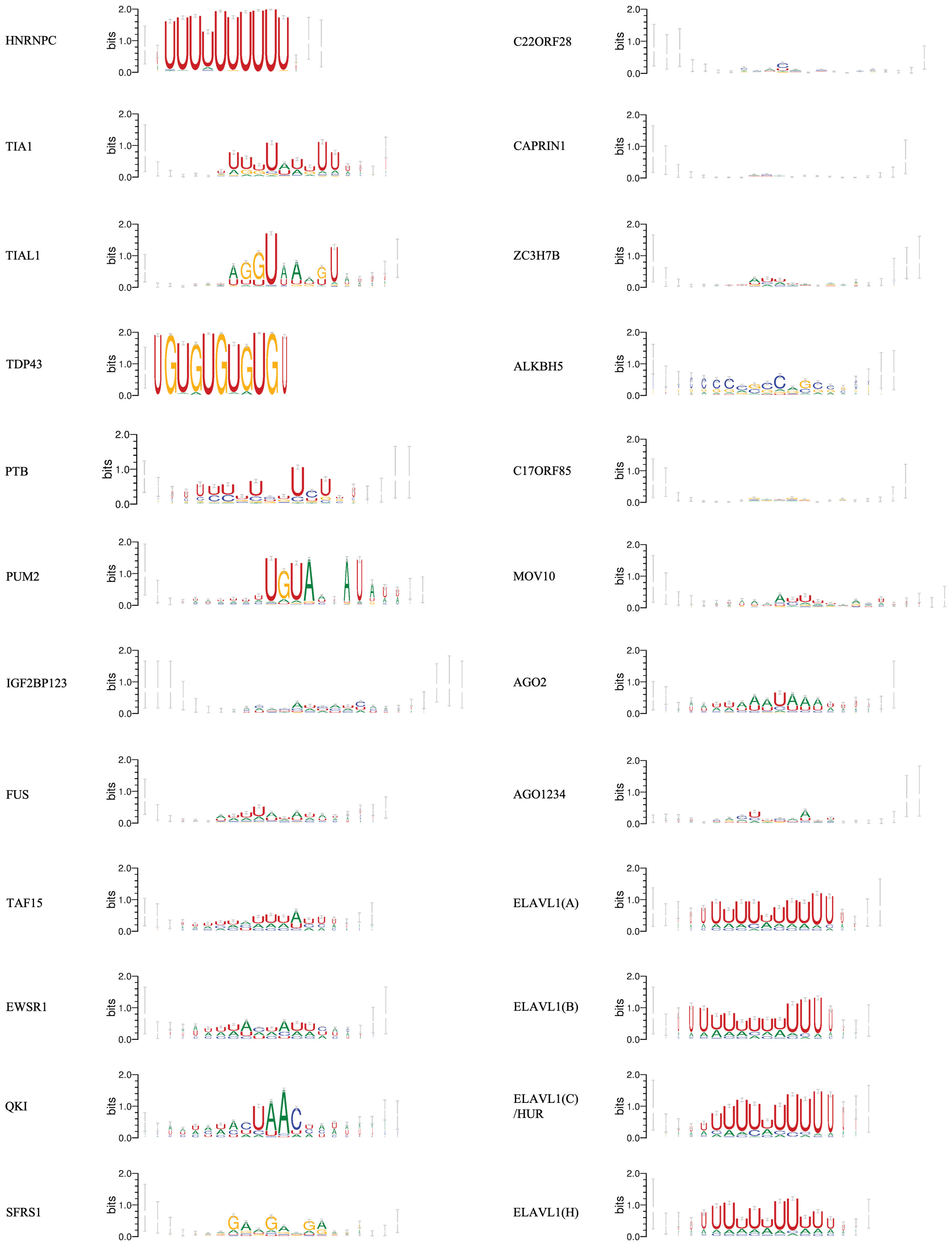


Figure S2: All sequence motifs extracted from our RPI-Net(CNN) model. Each one is aligned using 2000 10-mers.

### E Alternative sequence motifs discovered by MEME

| Protein | MEME viewpoints | MEME 10-mers |
|---|---|---|
| HNRNPC + Native BM | 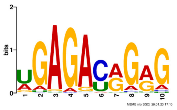 E-value: 2.9e-677 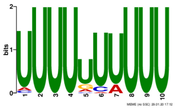 E-value: 5.8e-187 | Threshold: 0.6, 11782 10-mers<br>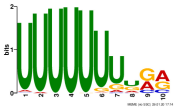 E-value: 2.5e-230 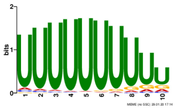 E-value: 6.4e-049 |
| TIA1 + Native BM | None | Threshold: 0.4, 10136 10-mers<br>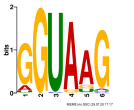 E-value: 1.2e-336 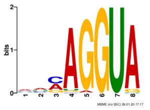 E-value: 9.2e-091 |
| TIAL1 + Native BM | 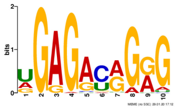 E-value: 6.3e-022 | Threshold: 0.3, 13123 10-mers<br>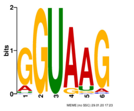 E-value: 1.2e-440 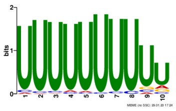 E-value: 4.0e-197 |
| TDP43 + Native BM | 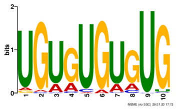 E-value: 5.0e-272 | Threshold: 0.3, 4540 10-mers<br>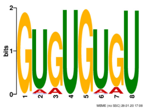 E-value: 1.9e-1645 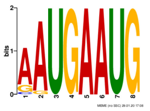 E-value: 8.7e-003 |
| PTB + Native BM | None | Threshold: 0.2, 875 10-mers<br>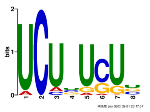 E-value: 2.9e-076 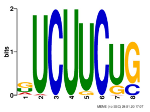 E-value: 1.3e-019 |
| PUM2 + Native BM | 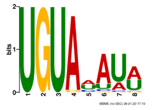 E-value: 2.0e-306 | Threshold: 0.1, 6635 10-mers<br>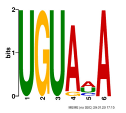 E-value: 6.6e-258 |
| IGF2BP123 + Native BM | None | Threshold: 0.1, 8558 10-mers<br>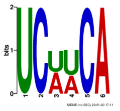 E-value: 2.8e-013 |
| FUS + Uniform BM | 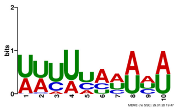 E-value: 9.7e-825 | Threshold: 0.3, 5390 10-mers<br>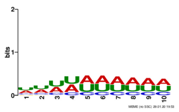 E-value: 1.1e-1056 |

| Protein | MEME viewpoints | MEME 10-mers |
|---|---|---|
| TAF15 + Uniform BM | 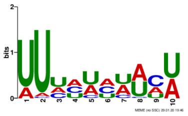<br>E-value: 2.2e-975 | Threshold: 0.3, 2662 10-mers<br>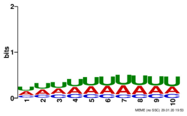<br>E-value: 6.4e-1152 |
| EWSR1 + Uniform BM | 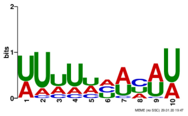<br>E-value: 2.2e-708 | Threshold: 0.1, 2945 10-mers<br>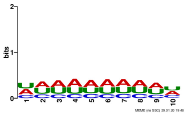<br>E-value: 5.9e-953 |
| QKI + Native BM | 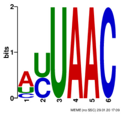<br>E-value: 9.9e-099 | Threshold: 0.1, 4375 10-mers<br>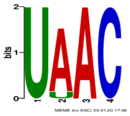<br>E-value: 6.3e-329 |
| SFRS1 + Native BM | 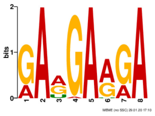<br>E-value: 1.1e-023 | Threshold: 0.2, 8103 10-mers<br>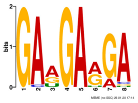 E-value: 6.4e-149<br>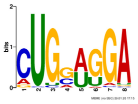 E-value: 3.8e-078<br>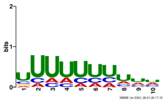 E-value: 1.8e-043 |
| C22ORF28 + Native BM | None | Threshold: 0.1, 4494 10-mers<br>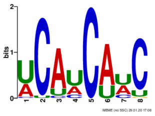<br>E-value: 2.2e-022 |
| CAPRIN1 + Native BM | None | Threshold: 0.3, 1961 10-mers<br>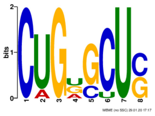<br>E-value: 2.2e-007 |
| ZC3H7B + Native BM | None | Threshold: 0.7, 2266 10-mers<br>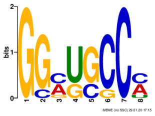<br>E-value: 8.0e-007 |
| ALKBH5 + Native BM | None | Threshold: 0.005, 332 10-mers<br>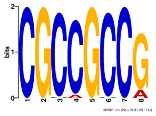<br>E-value:1.1e-034 |

| Protein | MEME all viewpoints | MEME 10-mers |
|---|---|---|
| C17ORF85 + Uniform BM | None | Threshold: 0.7, 811 10-mers<br>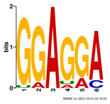 E-value: 8.5e-030<br>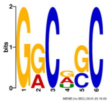 E-value: 1.2e-017<br>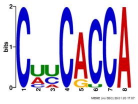 E-value: 9.6e-007 |
| MOV10 + Uniform BM | 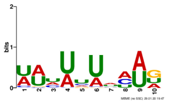<br>E-value: 4.1e-211 | Threshold: 0.5, 1302 10-mers<br>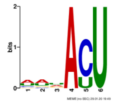 E-value: 5.2e-381<br>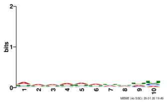 E-value: 1.6e-104 |
| AGO2 + Uniform BM | 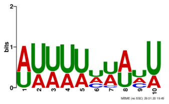<br>E-value: 7.2e-119 | Threshold: 0.3, 8130 10-mers<br>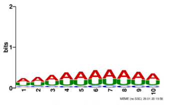 E-value: 1.0e-1016 |
| AGO1234 + Uniform BM | 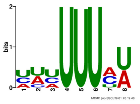<br>E-value: 2.5e-027 | Threshold: 0.3, 1403 10-mers<br>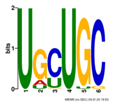 E-value: 1.4e-201<br>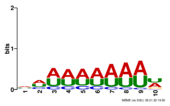 E-value: 9.2e-131<br>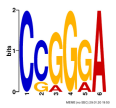 E-value: 6.0e-078<br>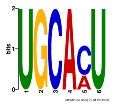 E-value: 7.9e-064<br>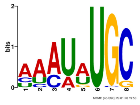 E-value: 2.8e-023 |
| ELAVL(A) + Uniform BM | 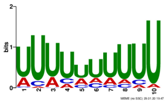<br>E-value: 4.2e-1484 | Threshold: 0.3, 18274 10-mers<br>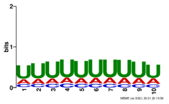 E-value: 1.5e-1853 |
| ELAVL(B) + Uniform BM | 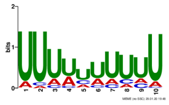<br>E-value: 8.1e-1362 | Threshold: 0.3, 11362 10-mers<br>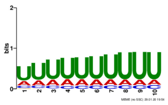 E-value: 3.3e-2250 |
| ELAVL(C) HUR + Uniform Model | 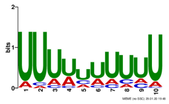<br>E-value: 7.2e-1660 | Threshold: 0.3, 13076 10-mers<br>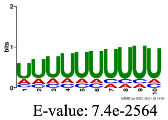 E-value: 7.4e-2564 |
| ELAVL(H) + Uniform BM | 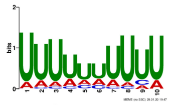<br>E-value: 1.4e-1624 | Threshold: 0.3, 1600 10-mers<br>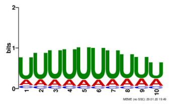 E-value: 2.1e-1907 |

Figure S3: In addition to the sequence motifs obtained with multiple sequence alignment, this figure presents some alternative motifs found by MEME on all the 10-mers whose integrated gradient scores are above some certain thresholds. MEME may discover some secondary motifs possibly indicating cooperative bindings, from the candidate 10-mers screened by our RPI-Net(CNN) models. We also ran MEME on the entire viewpoint regions from all positive sequences as a control for the quality of motifs; in many cases the motif found that way are not as significant as those found by limiting the analysis to high integrated gradients regions. "Native BM" indicates that we let MEME determine its own 0-th order background model from the input sequence, whereas "Uniform BM" indicate cases where MEME was required to use a uniform 0-order background model where A, C, G, U receive equal probabilities.

### F All RPI-Net(GNN) motifs

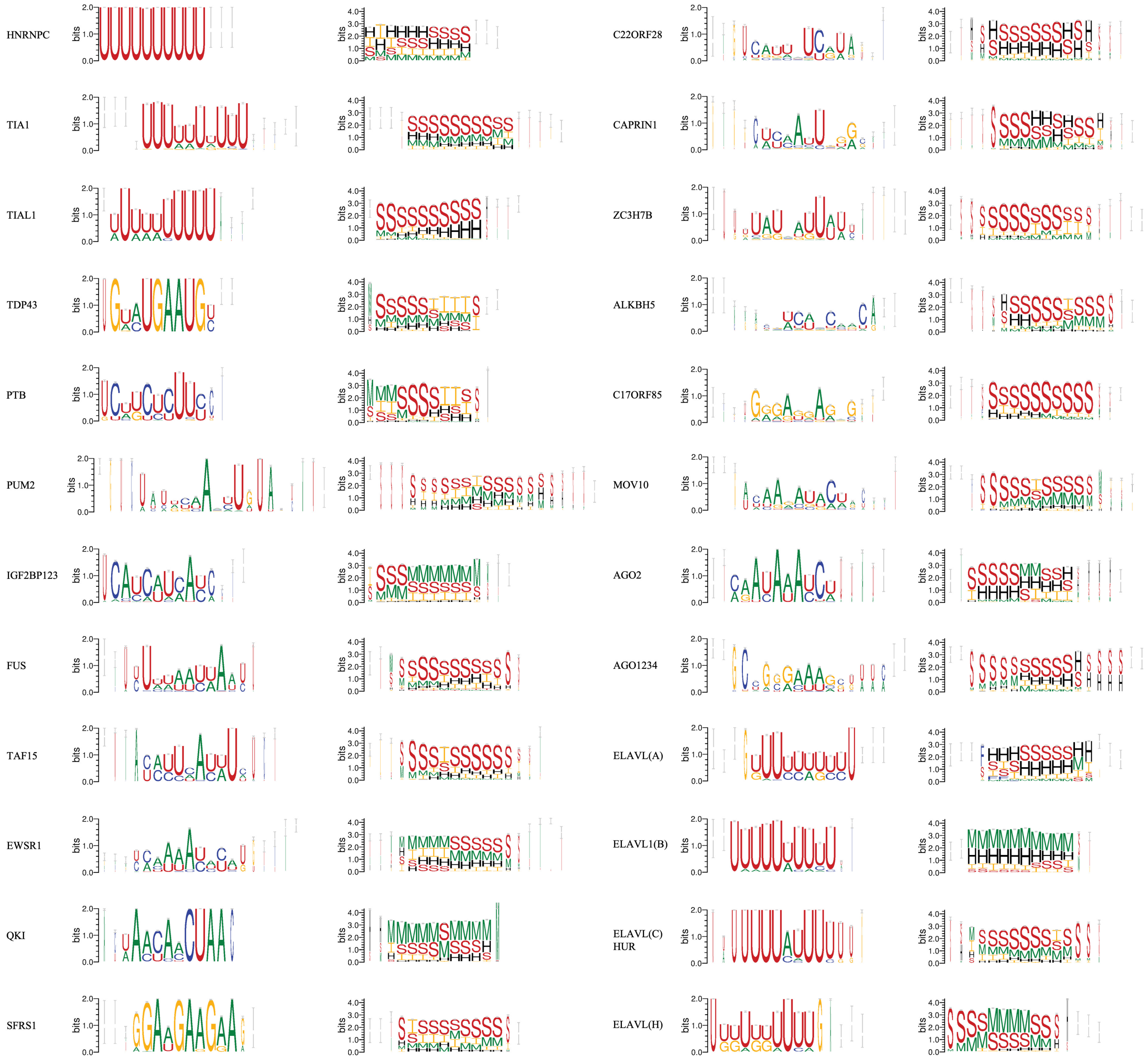


Figure S4: All sequence and secondary structural motifs extracted from our RPI-Net(GNN) model. Each one is aligned using 6000 10-mers.

#### G Secondary structure of RNA brings closer potential binding sites

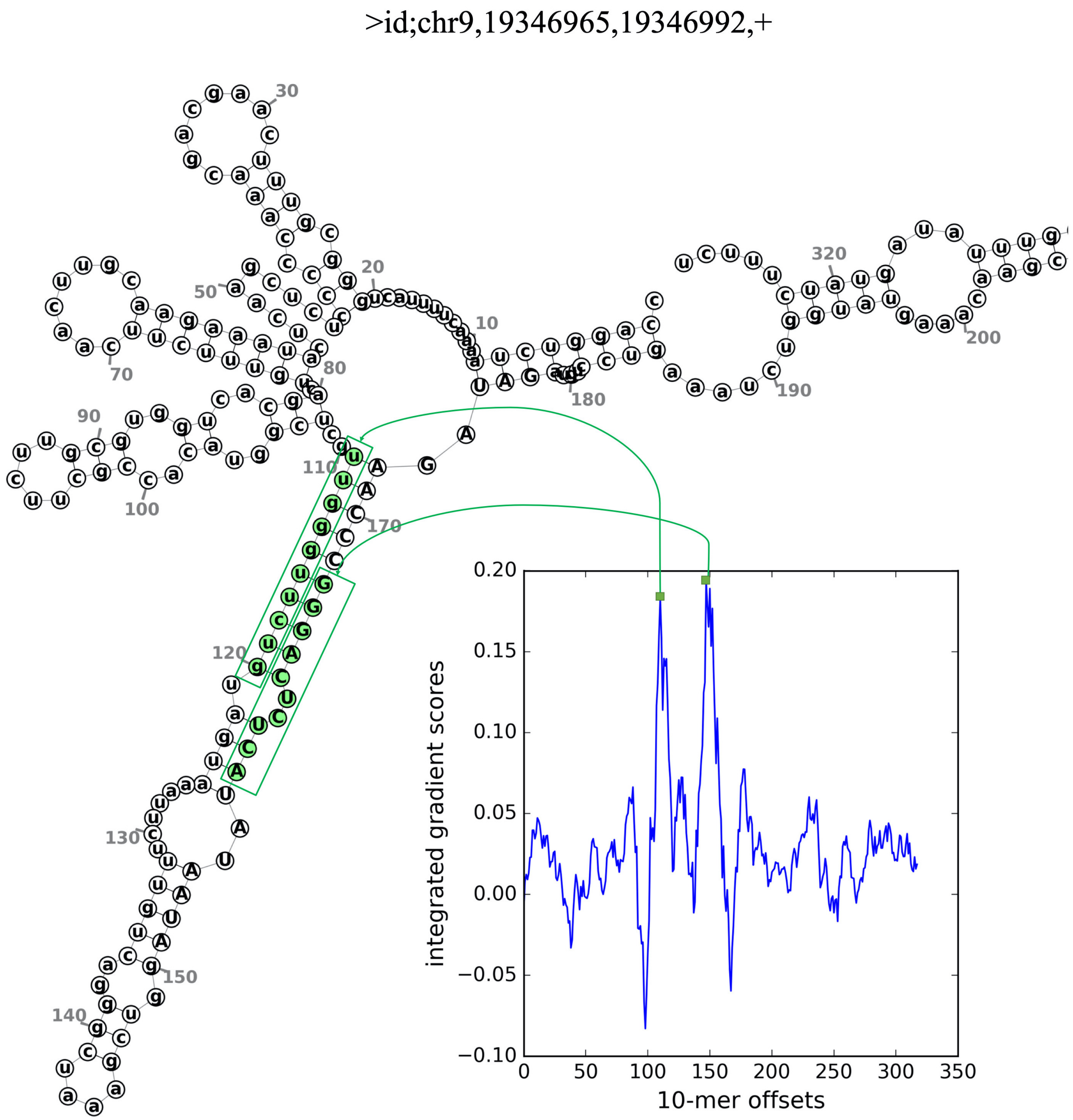


Figure S5: An example where the RNA secondary structure brings two potential binding sites into closer physical proximity. The plot in the lower right bottom shows the integrated gradient scores for all 10-mers along the RNA sequence, and the two distinct peaks indexed at 110 and 157 possibly indicate binding sites. The two 10-mers are mapped onto the minimum free energy structure and are shown in green.
